# Stress enhances hippocampal neuronal synchrony and prolongs sharp-wave ripples

**DOI:** 10.1101/2020.06.19.162487

**Authors:** Anupratap Tomar, Denis Polygalov, Sumantra Chattarji, Thomas McHugh

## Abstract

Chronic stress affects hippocampal function at multiple levels of neural organization. However, much of this understanding is derived from postmortem analyses of molecular, morphological, physiological and behavioral changes at fixed time points. Neural signatures of an ongoing stressful experience in the intact brain of awake animals and their links to later hippocampal dysfunction remain poorly understood. Here we used *in vivo* tetrode recordings to analyze the dynamic impact of 10 days of immobilization stress on neuronal activity in area CA1 of mice. Unexpectedly, there was a net decrease in pyramidal cell activity in stressed animals. Although these results suggest a lack of stress-induced hyperexcitability, more detailed analysis revealed that a greater fraction of spikes occurred specifically during sharp-wave ripples, resulting in an increase in neuronal synchrony. After repeated stress some of these alterations were visible during rest even in the absence of stress. These findings offer new insights into stress-induced alterations in ripple-spike interactions and mechanisms through which chronic stress may interfere with subsequent information processing.

## Introduction

The hippocampus is a medial temporal lobe structure that is crucial for encoding, updating and retrieving episodic memories (Eichenbaum, 2017; Tulving, 1985). Unfortunately, the same plasticity mechanisms that enable the hippocampus to perform these important functions, also make it vulnerable to damage caused by severe and repeated stress (Chattarji et al., 2015; McEwen et al., 2015). Stress-induced changes in the rodent hippocampus include shrinkage and debranching of pyramidal cell dendrites, (McEwen et al., 2015; Sousa et al., 2000), loss of dendritic spines (Magariños et al., 1997; Sandi et al., 2003) and alterations in synaptic plasticity mechanisms, including long-term potentiation (Alfarez et al., 2003; Shors et al., 1989). Together, these detrimental effects of stress at the cellular and synaptic levels are thought to contribute to impairments in hippocampal learning and memory. However, our current understanding is based primarily on postmortem analyses of stressed versus unstressed animals at fixed time points after the end of stress. The gradual and cumulative impact of stress on hippocampal function in the same animal over the course of repeated stress has not been explored in detail. Further, relatively little is known about how the intact, drug-free hippocampus is involved in the quick appraisal of an ongoing stressful situation (Cadle and Zoladz, 2015; Joëls, 2009).

These gaps in knowledge were addressed in several rodent studies that examined the effects of stress on theta-associated foraging behaviour (Kim et al., 2007; Park et al., 2015; Passecker et al., 2011; Tomar et al., 2015) during which hippocampal pyramidal ‘place’ cells (O’Keefe and Dostrovsky, 1971) form a cognitive map of the animals’ surroundings (O’Keefe and Nadel, 1978). Surprisingly, earlier reports found only modest changes in place cell properties between stressed and control subjects. However, in addition to theta-associated exploratory states, pyramidal cells also exhibit highly coordinated activity during off-line behavioural states such as rest and sleep (Buzsáki, 1989). These offline states are dominated by high-frequency (100-200 Hz) transients termed sharp-wave ripples (SPW-Rs) which have been shown to play key roles in learning and memory (Ego-Stengel and Wilson, 2010; Girardeau et al., 2009; Jadhav et al., 2012) and the planning of future actions (Joo and Frank, 2018; Pfeiffer and Foster, 2013; Schmidt and Redish, 2013). However, no information is available on the effects of either acute or chronic stress on SPW-R properties or their modulation of pyramidal cell activity. Further, if and how SPW-Rs are affected during stress, and if such effects differ between acute versus chronic stress remains to characterized. Hence, the current study aimed to address these outstanding issues. To this end, we examined hippocampal neural dynamics in mice on the first (acute) and last (chronic) day of a chronic immobilization stress (CIS) protocol. To characterize a neural signature of stress in CA1 we recorded local field potentials (LFPs) and neuronal activity during stress and compared it with activity recorded in an adjacent quiescence/rest state.

## Results

We examined the dynamics of CA1 pyramidal cell activity and ongoing LFPs on the first (acute) and last day (chronic) of 10-day chronic immobilization stress (CIS) (Fig. 1a), replicating a protocol that has been previously used to examine the effects of chronic stress on hippocampal dendrites (Vyas et al., 2002), spatial coding (Tomar et al., 2015), and behaviour (Ramirez et al., 2015; Suvrathan et al., 2010). Specifically, we examined the effects of stress at two time-points during CIS. First, we compared activity during the first episode of 2-hour stress (stress-state) to the preceding stress-free period (rest-state) on Day 1; this is referred to as the “acute” condition as this involves only a single exposure to stress. In other words, the effects of acute stress on the first day of CIS were analysed by comparing two conditions: i) acute-rest, and ii) acute-stress. Second, we carried out the same analyses on the last day of CIS, when the same animal has already experienced 9 exposures of the same stressor. Thus, we again compared activity during the 10^th^ episode of 2-hour stress (stress-state) to an adjacent stress-free period (rest-state). This is referred to as the “chronic” condition as it involves quantifying the cumulative effects of repeated stress over 10 days. Here again we compared two more conditions: iii) chronic-rest, and iv) chronic-stress, and together data from these four conditions will be presented in the following sections. The CIS protocol led to a gradual decrease in body weight (acute, 28.82 ± 0.97 vs chronic, 26.65 ± 0.69, N = 4 mice, paired t-test: t = 3.8411, p = 0.031) (supplementary Fig. 1a), confirming the efficacy of this chronic stress paradigm as it is consistent with previous reports (Vyas et al., 2002). Lesions in the stratum pyramidale confirmed that recordings were made from area CA1 (Fig. 1b).

**Fig. 1.**
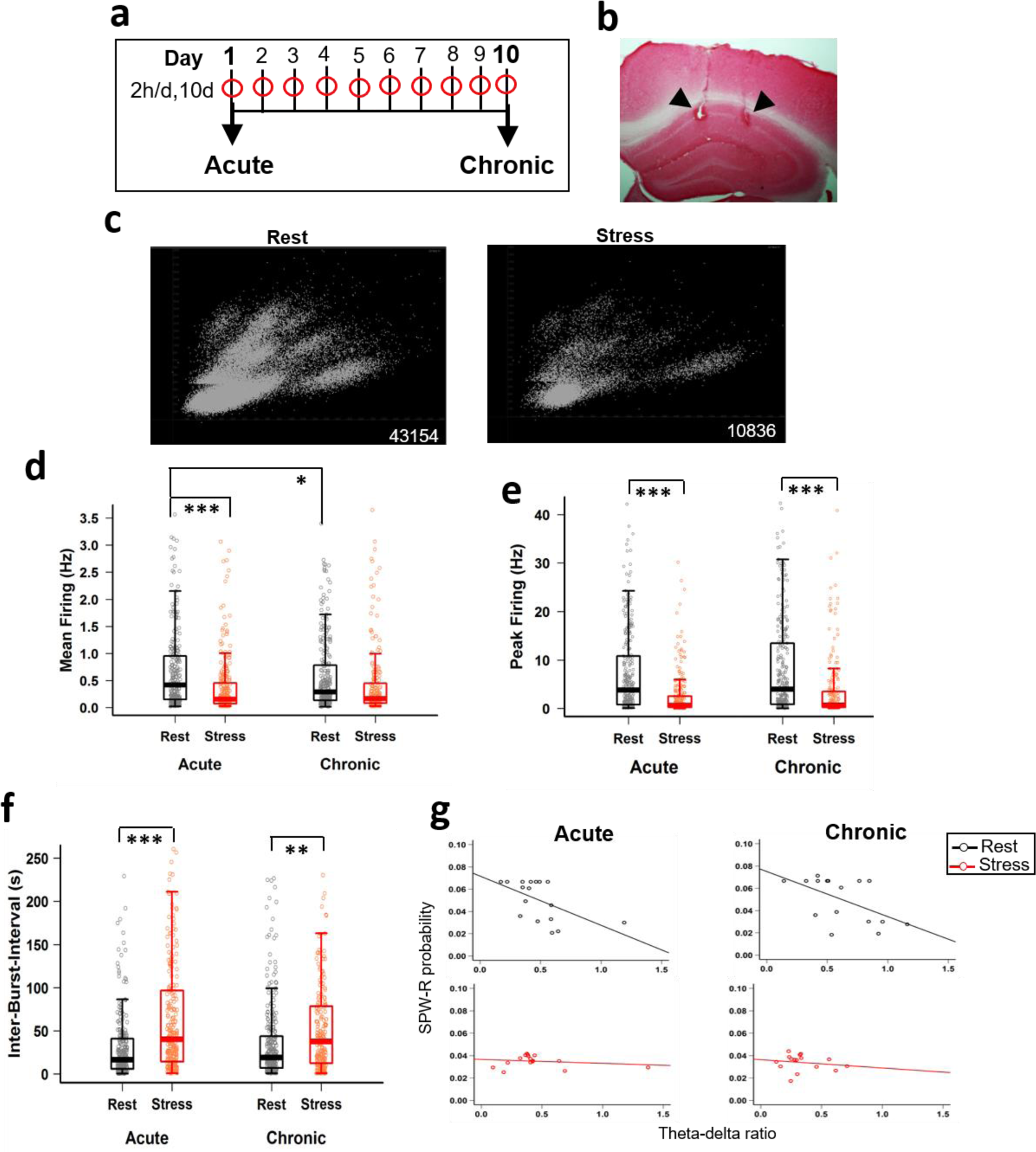
CA1 pyramidal cell activity is altered during stress. **a** Schematic representation of the chronic immobilization stress (CIS) protocol that mice received every day for 10 consecutive days. **b** Coronal section of the hippocampus showing the tetrode locations (black arrows) at the CA1 pyramidal layer. **c** Time matched (30 min) representative examples of unsorted spikes recorded during rest-state (left) and stress-state (right). Numbers next to each cluster depict total number of spikes recorded during that state. **d** Mean firing rate between rest-state and stress-state (two-way ANOVA: behaviour-state, F(1, 1074) = 29.611, p = 6.54×10^−8^; day, F(1, 1074) = 0.908, p = 0.341; behaviour-state x day interaction, F(1,1074) = 9.118, p = 0.0026; Tukey’s HSD: acute-rest vs acute-stress, p = 1.32×10^−8^; acute-rest vs chronic-rest, p=0.017; chronic-rest vs chronic-stress, p = 0.42). **e** Peak firing rate between rest-state and stress-state (two-way ANOVA: behaviour-state, F(1, 1074) = 54.234, p = 3.54×10^−13^; day, F(1, 1074) = 4.014, p = 0.0454; behaviour-state x day interaction, F(1, 1074) = 0.005, p = 0.945; Tukey’s HSD: acute-rest vs acute-stress, p = 7.0×10^−7^; chronic-rest, vs chronic-stress, p = 2.5×10^−6^). **f** Inter-burst-interval between rest-state and stress-state (two-way ANOVA: behaviour-state, F(1, 1066) = 47.294, p = 1.04×10^−11^; day, F(1, 1066) = 0.749, p = 0.387; behaviour-state x day interaction, F(1, 1066) = 3.685, p = 0.055; Tukey’s HSD: acute-rest vs acute-stress, 2.2×10^−9^; chronic-rest vs chronic-stress, p = 0.005). **g** Correlation between theta/delta ratio and SPW-Rs differs between behaviour-states on the first day (left: acute-rest, slope = -0.044, R = -0.56, p = 0.02; acute-stress, slope = 0.00, R = -0.19, p =0.47) and last day (right: chronic-rest, slope = -0.04, R = -0.54, p=0.029; chronic-stress, slope = -0.01, R = -0.17, p = 0.52) of CIS. All box plots represent interquartile range (IQR, 25^th^-75^th^ percentiles), median is the thick line in the box and whiskers extend to 1.5 times the IQR. Circles depict rest-state (black) stress-state (red). * p<0.05, ** p < 0.01, *** p < 0.001. acute-rest: n = 288 cells, N = 17 mice, acute-stress: n = 282 cells, N = 16 mice, chronic rest: n = 282 cells, N = 16 mice, chronic stress: n = 226 cells, N = 16 mice).

### CA1 pyramidal cell spiking differs between stress-state and rest-state

A long-held view on the detrimental effects of stress centers on the idea that severe and repeated stress leads to hippocampal hyperactivity, which in extreme cases may cause excitotoxic damage. For instance, both the glucocorticoid cascade (Sapolsky, 1996) and synaptic saturation hypotheses of stress (Cadle and Zoladz, 2015; Diamond et al., 2004) suggest that enhanced calcium and glutamate release during stress alter subsequent synaptic plasticity and mnemonic processes. The implicit assumption underlying both these hypotheses is that hippocampal neuronal networks undergo hyperexcitability during stress though no study has directly tested this possibility in the intact brain *in vivo*. Surprisingly, we found hippocampal multiunit activity to be suppressed during stress (Fig. 1c and supplementary Fig. 1b, c). Comparison of the average firing rate of CA1 pyramidal cells between stress and rest across days revealed a significant decrease during the stress-state (Fig 1d) with the difference being most pronounced during acute stress on the first day (acute-rest, 0.80 ± 0.06 Hz vs acute-stress, 0.38 ± 0.03 Hz, p = 1.32×10^−8^, Tukey’s HSD). Following repeated stress, the mean firing rate continued to be low not only during the last session of chronic stress, but also during the adjacent rest period, resulting in a significant reduction of firing rates was both the rest and stress periods on Day 10, indicative of carryover of the stress phenotype into non-stress states as well (acute-rest, 0.80 ± 0.06 Hz, vs chronic-rest, 0.59 ± 0.04 Hz, p = 0.017; chronic-rest, 0.59 ± 0.04, vs chronic-stress, 0.48 ± 0.05, p = 0.425, Tukey’s HSD). Similarly, we observed a significant lowering of peak firing rates (Hz) (Fig. 1e) on both the first day (acute-rest, 7.77 ± 0.59 Hz, vs acute-stress, 3.37 ± 0.49 Hz, p = 7.41×10^−7^, Tukey’s HSD) and the last day (chronic-rest, 8.80 ± 0.69 Hz vs chronic-stress, 4.31 ± 0.62 Hz, p = 2.53×10^−6^, Tukey’s HSD) of CIS. While stress did not alter the length (ms) of bursting activity, the stress-state was associated with longer inter-burst-intervals (Fig. 1f), indicating that time gaps (s) between burst activity were lengthened, consistent with the overall decrease in activity (acute-rest, 33.29 ± 3.33 vs acute-stress, 78.69 ± 6.42, p = 2.2×10^−9^; chronic-rest, 40.41 ± 3.68 vs chronic-stress, 65.33 ± 6.82, p = 5.35 ×10^−3^, Tukey’s HSD). Taken together, these data demonstrate that stress decreases the activity of CA1 pyramidal cells.

### LFP profile and SPW-R properties differ between the stress-state and rest-state

Considering stress did not cause a straightforward increase in CA1 firing rates, we next focused our attention on other measures of hippocampal neural dynamics. Hippocampal behavioral states display characteristic patterns of LFP activity (Buzsáki, 1989) wherein rest and immobility states are defined by a low ratio of LFP power in the theta (6-12 Hz) and delta (1-4 Hz) bands, along with periodic high-frequency sharp-wave ripples (SPW-Rs, 150-200 Hz) (Grosmark et al., 2012). Consistent with earlier reports, we observed a negative correlation between the theta/delta ratio and SPW-R probability during the unstressed rest-state. But, this relationship was disrupted during both acute (Fig. 1g, left; acute-rest, R = -0.56, p = 0.02, vs acute-stress, R = -0.19, p = 0.47) and chronic stress (Fig. 1a, right; chronic-rest, R = -0.54, p = 0.029 vs chronic-stress, R = -0.17, p = 0.52), raising the possibility that CIS induces a unique physiological state.

The above findings underscore the need to examine in greater detail the effects of stress on SPW-Rs. Moreover, there is growing evidence for SPW-Rs playing an important role in memory consolidation, which is affected by stress (Fernández-Ruiz et al., 2019; Nakashiba et al., 2009). Hence, we next analyzed SPW-R properties in both the early and late stages of chronic stress. While no differences were observed in the intrinsic oscillation frequency (supplementary Fig. 2a), the average SPW-R duration was significantly longer during stress at both acute and chronic time points (Fig. 2b; acute-rest, 104.94 ± 3.29 ms vs acute-stress, 137.05 ± 4.65 ms, p= 3.86×10^−5^; chronic-rest, 107.54 ± 2.44 ms vs chronic-stress, 130.35 ± 6.996 ms, p = 0.0055, Tukey’s HSD). The population distribution of SPW-R duration (Fig. 2c) clearly showed a greater fraction of long-duration ripples during both acute and chronic stress (acute-rest (median = 90.93) vs acute-stress (median = 122.27), p < 2.22×10^−16^; chronic-rest (median = 92.16) vs chronic-stress (median = 105.68), p < 2.22×10^−16^, Kolmogorov-Smirnov test). Further, the increase in SPW-Rs duration was characterized by a rapid onset and sustained increase following stress initiation (Fig. 2d).

**Fig. 2.**
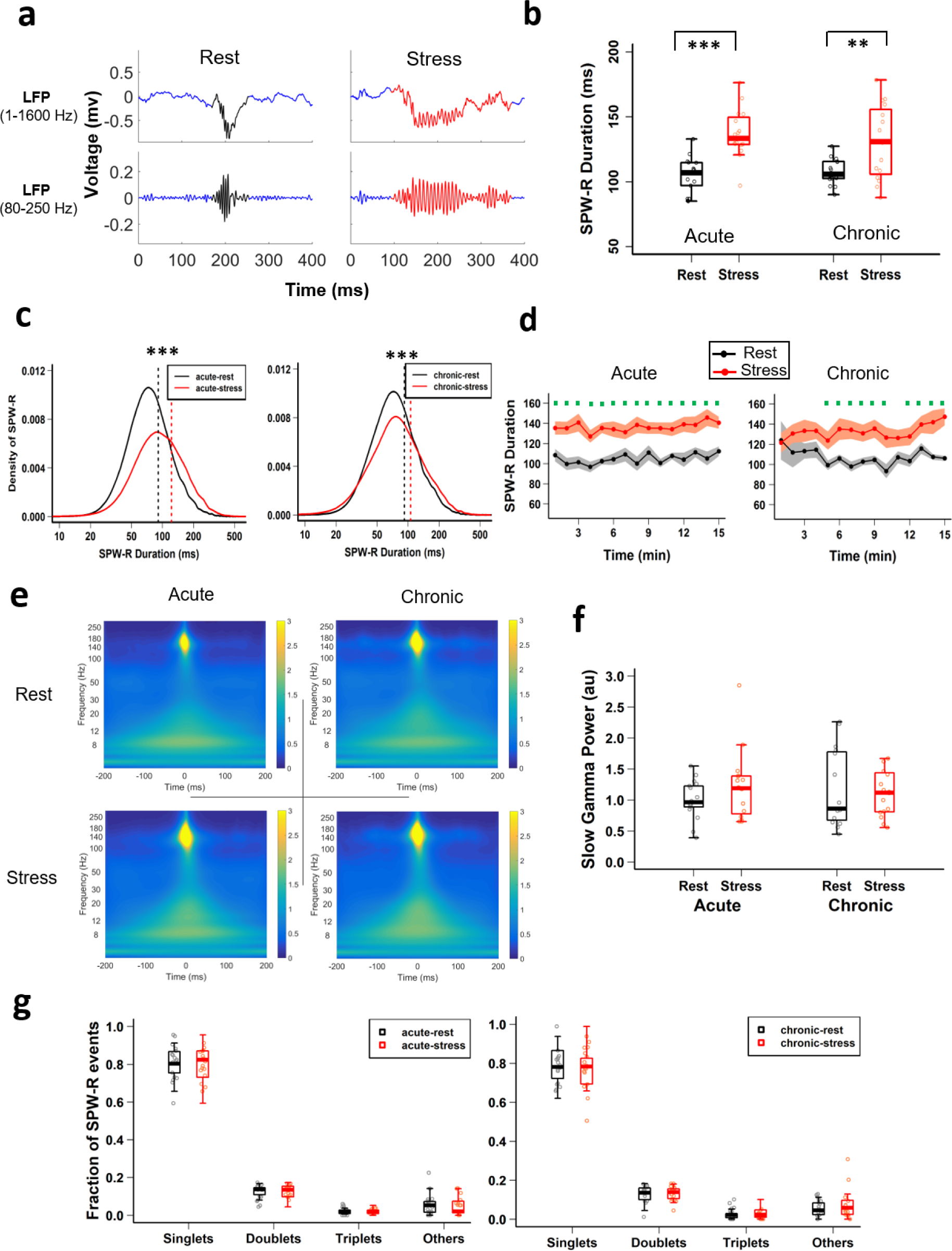
CA1 SPW-R duration differs between rest and stress states. **a** Representative examples of local field potential (LFP) for non-filtered (top) and filtered (bottom) SPW-R events recorded during rest-state (left) and stress-state (right) from CA1 *stratum pyramidale*. **b** SPW-R duration differs between rest-state and stress-state; (two-way ANOVA: behaviour-state, F(1, 61) = 35.233, p = 1.5×10^−7^; day, F(1, 61) = 0.114, p = 0.737; behaviour-state x day, F(1, 61) = 1.002, p = 0.321; Tukey’s HSD: acute-rest vs acute-stress, p = 3.86×10^−5^, chronic-rest vs chronic-stress, p = 0.0055). **c** Distributions of the duration of SPW-Rs differs between behaviour states on first day (left: acute-rest vs acute-stress, Kolmogorov-Smirnov test, D = 0.236, p < 2.22×10^−16^) and last day (right: chronic-rest vs chronic-stress, Kolmogorov-Smirnov test, D = 0.113, p < 2.22×10^−16^) showed a rightward shift. **d** Temporal dynamics of averaged SPW-R duration (1-min bins) differ between behaviour states on the first day (left: LMMs; behaviour-state, F(1, 447) = 249. 26, p < 2.22×10^−16^; time, F(14, 447) = 0.703, p = 0.772; behaviour-state x time, F(14, 447) = 0.456, p = 0.955). Similarly, on the last day, SPW-R duration increased during stress (right: LMMs; behaviour-state, F(1, 412) = 68.75, p = 1.74×10^−15^; time: F(14, 412) = 1.05, p = 0.404; behaviour-state x time, F(14, 412) = 1.226, p = 0.253). **e** Averaged peri-SPW-R wavelet spectrograms during acute-rest (top left), acute-stress (bottom left), chronic-rest (top right) and chronic-stress (bottom right). **f** Low gamma (17-40Hz) power during SPW-R events does not differ between behaviour-states (two-way ANOVA: behaviour-state, F(1, 61) = 1. 31, p = 0.26; day, F(1, 61) = 0.55, p = 0.46; behaviour-state x day, F(1, 61) = 0.07, p = 0.79). **g** Fraction of different types of ripple bursts during acute stress (two-way ANOVA: behaviour-state, F(1, 124) = .00, p = 1; category, F(3, 124) = 1521.05, p < 2.22×10^−16^; behaviour-state x category, F(3, 124) = 0.22, p = 0.88; Tukey’s HSD: acute-rest vs acute-stress: singlets, p = 0.999; doublets, p = 0.999; triplets, p = 0.999; others, p = 0.999) and chronic stress (two-way ANOVA: behaviour-state, F(1, 120) = .00, p = 1; category, F(3, 120) = 981.21, p < 2.22×10^−16^; behaviour-state x category, F(3, 124) = 0.169, p = 0.917; Tukey’s HSD: chronic-rest vs chronic-stress: singlets, p= 1.0; doublets, p=0.999; triplets, p=0.999; others, p=0.999). All box plots represent median and 25^th^-75^th^ percentiles, with whiskers extending to the extreme data points. Circles depict rest-state (black) stress-state (red). * p < 0.05, ** p < 0.01, *** p < 0.001. Acute-rest: n = 7220 SPW-Rs, N = 17 mice; acute-stress: n = 9851 SPW-Rs, N = 16 mice; chronic rest: n = 7041 SPW-Rs, N = 16 mice; chronic stress: n = 8978 SPW-Rs, N = 16 mice).

Recent work has suggested that longer SPW-Rs could result from the merging or concatenation of single ripple events, which is reflected as an increase in power in the underlying gamma oscillation (Oliva et al., 2018). To examine this possibility we next measured the low-gamma power (17-40 Hz) during ripple events, but found no differences between stress and rest during both the acute (Day 1) and chronic (Day 10) time points (Fig. 2e and f), suggesting absence of excessive merging of SPW-Rs. We also examined if stress leads to increased ripple bursts, a phenomenon ascribed to input from the entorhinal cortex (Yamamoto and Tonegawa, 2017). The proportion of ripples occurring as singlets (Fig. 2g) was similar between rest and stress at both the acute and chronic time points (singlets, acute-rest, 0.81 ± 0.02 vs acute-stress, 0.80 ± 0.02, p= 0.999; chronic-rest, 0.78 ± 0.02 vs chronic-stress, 0.78 ± 0.03, p= 1.0, Tukey’s HSD) again suggesting that the observed SPW-Rs were longer single events.

We next compared the normalized amplitudes (supplementary Fig. 2b) of SPW-Rs and found them to be significantly larger during both acute and chronic stress sessions (acute-rest, 4.98 ± 0.21 vs acute-stress, 6.22 ± 0.30, p = 0.013; chronic-rest, 5.14 ± 0.20 vs chronic-stress, 6.50 ± 0.37, p = 0.006, Tukey’s HSD). Once again, the increase in SPW-R amplitude showed a rapid onset and sustained increase following the beginning of the 2-hour stress on both the first and last days (supplementary Fig. 2c). These data demonstrated that stress induction quickly alters basic SPW-R properties in hippocampal area CA1.

### Stress alters excitability and participation of CA1 pyramidal cells during SPW-Rs

In addition to its hypothesized role as a biomarker of cognition, SPW-Rs have also been described as a “built in” physiological sensor of hippocampal excitability because of network wide activation of hippocampal neurons which accompanies their occurrence (Buzsáki, 2015). Thus, next we searched for evidence of hyperexcitability during both acute and chronic stress by comparing the pyramidal cell firing patterns during SPW-R events between rest and stress states. In agreement with suppressed average firing rates reported above, multiunit activity was muted during SPW-Rs across both acute and chronic stress sessions, though the magnitude of difference was larger during chronic stress (Fig. 3a). Neither acute nor chronic stress altered the relationship between CA1 pyramidal cell firing in SPW-Rs and their overall mean firing rate (supplementary Fig. 3a: acute-rest, R^2^ = 0.56, p < 2.22×10^−16^; acute-stress, R^2^ = 0.47, p < 2.22x 10^−16^, likelihood ratio test, p = 0.80; chronic-rest, R^2^ = 0.44, p < 2.22×10^−16^; chronic-stress, R^2^ = 0.27, p < 2.22×10^−16^, likelihood ratio test, p = 0.105). Further, similar to overall activity, we observed a decrease in pyramidal cell firing rate (Hz) inside SPW-Rs during acute stress (Fig 3b acute-rest, 2.20 ± 0.19 Hz vs acute-stress, 1.38 ± 0.12 Hz, p = 1.78×10^−4^, Tukey’s HSD, while following chronic stress, mean firing remained low both during rest and stress on the last day (acute-rest, 2.20 ± 0.19 Hz vs chronic-rest, 1.64 ± 0.11 Hz, p = 0.024, chronic-rest, 1.64 ± 0.11 Hz vs chronic-stress, 1.43 ± 0.11 Hz, p = 0.749, Tukey’s HSD). Further, the impact of chronic stress on firing in SPW-Rs during the rest-state was also evident in the cumulative distribution plots (supplementary Fig 3b), as the rest-state on the last day showed a leftward shift (acute-rest vs chronic-rest, p = 0.013, Tukey’s HSD). Firing outside of SPW-Rs changed in a similar fashion and decreased during acute-stress and chronic-rest states (supplementary Fig. 3c &d). Interestingly, despite this overall drop in activity, during stress-states pyramidal cells discharged a much larger percentage of their total spikes inside of SPW-Rs (Fig. 3c). The fraction of spikes inside SPW-Rs during stress almost doubled on the first day (acute-rest, 14.49 ± 0.85% vs acute-stress, 26.19 ± 1.21% p = 1.37×10^−12^, Tukey’s HSD), and remained elevated on the last day (chronic-rest, 15.93 ± 1.05% vs chronic-stress, 25.0.± 1.45%, p = 3.62×10^−7^,Tukey’s HSD) of CIS.

**Fig. 3.**
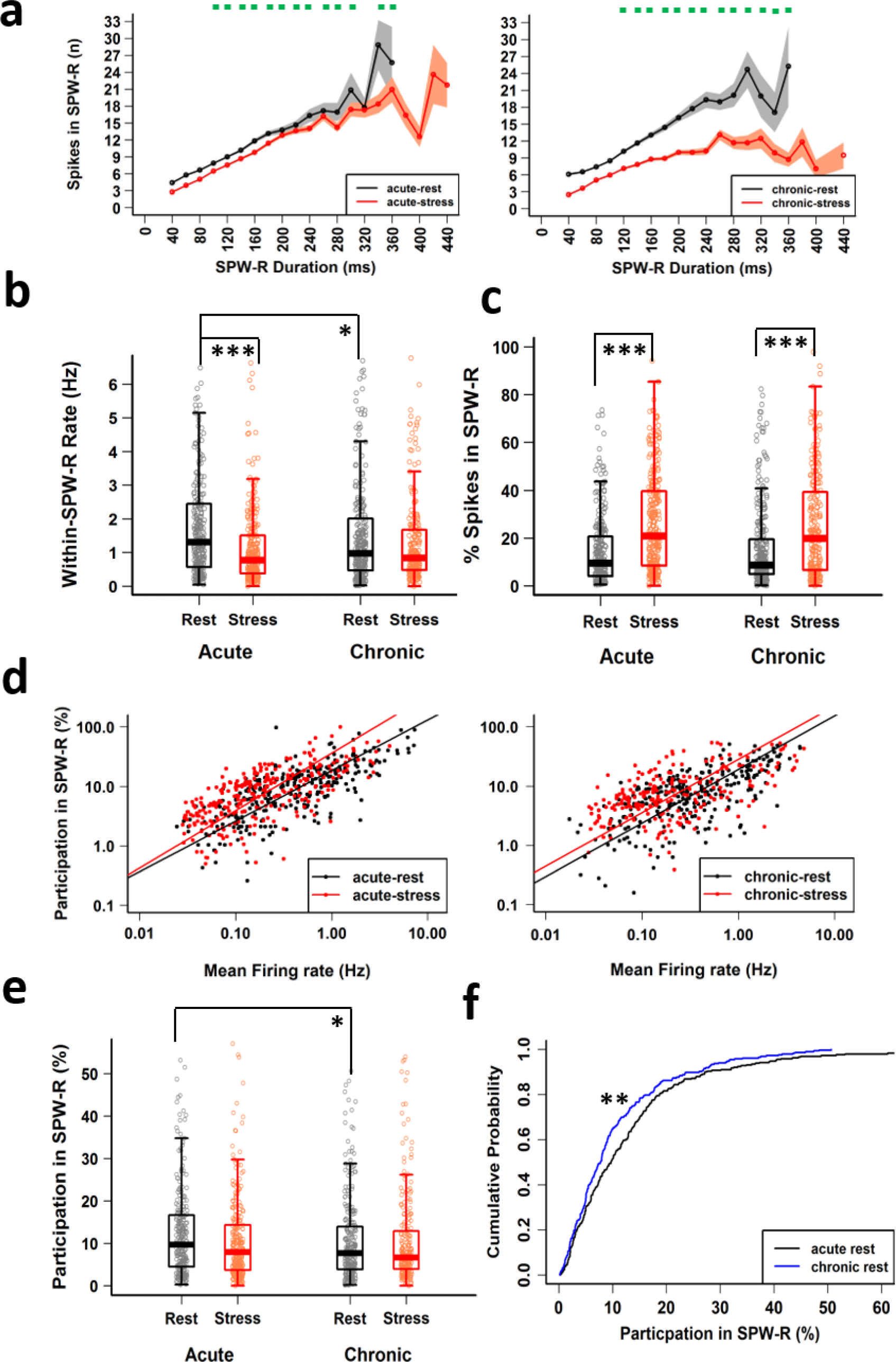
Altered CA1 pyramidal activation in SPW-Rs during stress-state. **a** The relationship between SPW-R duration (20-ms bins) and multiunit activity during SPW-Rs for first (left) and last (right) day of CIS (LMMs: behaviour-state, F(1, 16896) = 0.2325, p = 0.63; duration, F(16, 16896) = 715.93, p < 2.22×10^−16^; behaviour-state x duration, F(16, 16896) = 7.615, p < 2.22×10^−16^) and the last day (LMMs: behaviour-state, F(1, 15877) = 1.877, p = 0.17; duration, F(16, 15877) = 525.5, p < 2.22×10^−16^; behaviour-state x day, F(16, 15877) = 36.435, p < 2.22×10^−16^) of CIS). Green dots on top of graph indicate significant differences between behaviour-states. **b** Within SPW-R average firing rate differs between behaviour-states (two-way ANOVA: behaviour-state, F(1, 1074) = 13.981, p = 1.9×10^−4^; day, F(1, 1074) = 2.892, p = 0.089; behaviour-state x day, F(1, 1074) = 4.564, p = 0.033; acute-rest vs acute-stress, p = 1.78×10^−4^; chronic-rest vs chronic-stress, p = 0.749; acute-rest vs chronic-rest, p = 0.024). **c** Percentage of spikes discharged by CA1 pyramidal cells in SPW-R differs between behaviour-states (two-way ANOVA: behaviour-state, F(1, 1074) = 84.83, p < 2.22×10^−16^; day F(1, 1074) = 0.079, p = 0.778; behaviour-state x day, F(1, 1074) = 1.35, p = 0.25; Tukey’s HSD: acute-rest vs acute-stress, p = 1.4×10^−12^; chronic-rest vs chronic-stress, p = 3.62×10^−7^). **d** Dependency of pyramidal cell participation in SPW-Rs on their mean firing rate on the first day (acute-rest: slope = 0.848, R^2^ = 0.55, p < 2.22×10^−16^; acute-stress: slope = 0.96, R^2^ = 0.40, p < 2.22×10^−16^; likelihood ratio test (df = 1) = 4.318, p = 0.038) and the last day (chronic-rest: slope = 0.903, R^2^ = 0.42, p < 2.22×10^−16^; chronic-stress: slope=0.898, R^2^=0.21, p=2.97×10^−13^; likelihood ratio test (df = 1) = 0.006, p = 0.94) of CIS. **e** Pyramidal cell participation in SPW-Rs differs with the progression of stress protocol (two-way ANOVA: behaviour-state, F(1, 1074) = 1.686, p = 0.194; day, F(1, 1074) = 5.695, p = 0.017; behaviour-state x day, F(1, 1074) = 2.19, p = 0.139; Tukey’s HSD: acute-rest vs acute-stress, p = 0.204; chronic-rest vs chronic-stress, p = 0.998; acute-rest vs chronic rest, p = 0.027). **f** Cumulative distribution of pyramidal cell participation in SPW-Rs differs between rest-states on the first day (black) and last day (blue) of CIS (Kolmogorov-Smirnov test: D = 0.142, p = 0.006). All box plots, represent interquartile range (IQR, 25^th^-75^th^ percentiles), median is the thick line in the box and whiskers extend to 1.5 times the IQR. * p < 0.05, ** p < 0.01, *** p < 0.001. Acute-rest: n = 288 cells, N = 17 mice; acute-stress: n = 282 cells, N = 16 mice; chronic rest: n = 282 cells, N = 16 mice; chronic stress: n = 226 cells, N = 16 mice).

We next assessed the effects of stress on the participation of pyramidal cells in SPW-Rs events. In agreement with a previous reports (Fernández-Ruiz et al., 2019; O’Keefe and Nadel, 1978), the baseline firing rate of pyramidal cells displayed a positive relationship with the cell’s participation probability in SPW-Rs during both acute and chronic stress, though slight but significant differences were observed during acute stress (Fig. 3d: acute rest, R^2^ = 0.55, p < 2.22×10^−16^; acute-stress, R^2^ = 0.40, p < 2.22×10^−16^; likelihood ratio test, p = 0.038; chronic-rest, R^2^ = 0.42, p < 2.22×10^−16^; chronic-stress, R^2^ = 0.21, p = 2.97×10^−13^, likelihood ratio test, p = 0.94). Further, the extent of pyramidal cell participation in SPW-Rs varied widely (Fig. 3e;(Grosmark et al., 2012; Ylinen et al., 1995). However, a comparison across sessions and days revealed a significant effect of day (Fig. 3e); while pyramidal cell participation was similar on the first day (acute-rest, 13.39 ± 0.84% vs acute-stress, 11.41 ± 0.73%, p = 0.204, Tukey’s HSD), the chronic rest-state showed a significantly lower participation than that of the acute rest-state (acute-rest, 13.39 ± 0.84% vs chronic-rest, 10.57 ± 0.58%, p = 0.027, Tukey’s HSD), suggesting that chronic stress suppresses pyramidal cell participation in SPW-Rs. No further decrease was observed between behaviour states on the last day (chronic-rest, 10.57 ± 0.60% vs chronic-stress, 10.77 ± 0.73%, p = 0.997, Tukey’s HSD) of CIS. Cumulative distribution plots further confirmed the impact of chronic stress on SPW-R participation during the rest state (Fig. 3f: acute-rest vs chronic-rest, p = 0.006, Kolmogorov-Smirnov test). Overall, these data demonstrated that despite suppressed firing rates, SPW-R-specific activation of CA1 pyramidal cells was enhanced during both acute and chronic stress. However, with progression of the CIS protocol, pyramidal cells participated in fewer SPW-Rs during the rest period adjacent to the stress exposure.

### Co-firing of CA1 pyramidal cells is altered by stress

Hippocampal neuronal synchrony peaks during SPW-Rs (Buzsáki et al., 1992; Csicsvari et al., 1999a; Wilson and McNaughton, 1994) and this has been suggested as a key mechanism in hippocampal mnemonic function (Buzsáki and Mizuseki, 2014; Cheng and Frank, 2008). Our finding that a greater fraction of pyramidal spiking occurs inside SPW-Rs, raised the possibility of enhanced synchrony of pyramidal cell firing during stress. However, the overall decrease in firing rates, and reduced SPW-R participation by pyramidal cells with repeated stress, suggested the contrary. To examine this, we quantified the co-activation of pairs of pyramidal cells exhibiting a positive correlation in their firing patterns in a given session and observed a significant elevation during stress periods (Fig. 4a). Co-activity Z-scores were significantly higher during acute (acute-rest, 1.228 ± 0.027 vs acute-stress, 1.495 ± 0.031, p = 1.2×10^−8^, Tukey’s HSD) as well as chronic stress (chronic-rest, 1.388 ± 0.03, vs chronic-stress, 1.622 ± 0.043, p = 3.8×10^−6^, Tukey’s HSD). Further, the rest-state on the last day showed significantly larger co-activation values as compared to the first day (acute rest vs chronic rest, p = 0.0016, Tukey’s HSD). Finally, to compare the coordinated activity more directly, we calculated co-activity scores of cell-pairs with positive values simultaneously recorded both rest and stress states (Fig. 4b). This analysis revealed a positive relationship between cell pairs during both rest and stress states on the first day (acute: R^2^ = 0.022, p = 0.033), which grew stronger with further episodes of stress over time (chronic: R^2^ = 0.019, p = 0.076) and significantly differed between days (p = 1.28×10^−7^, likelihood ratio test) suggesting that chronic stress led to reorganization of network activity even during non-stress periods.

**Fig. 4.**
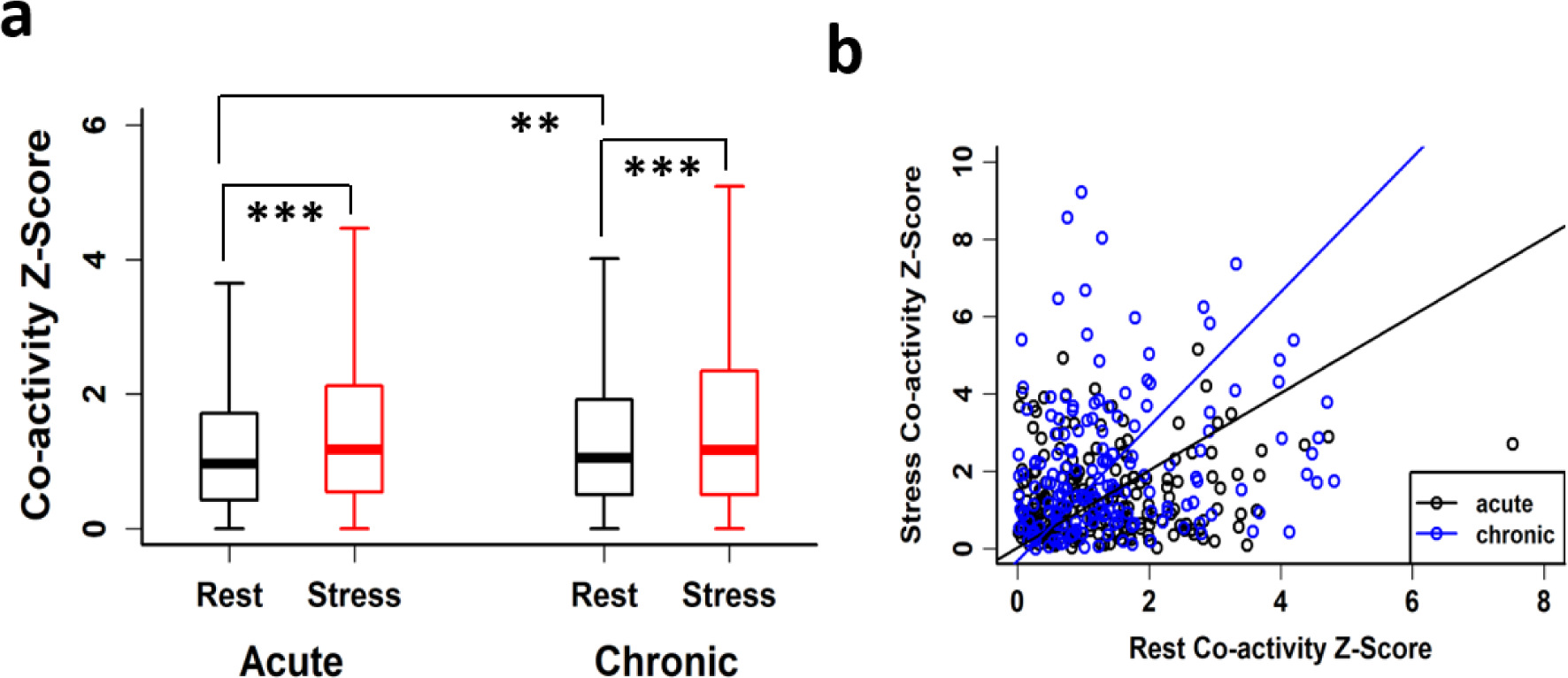
Altered CA1 pyramidal cell coactivation in SPW-Rs during stress-state. **a** Co-activity Z-scores of pyramidal cells during SPW-Rs differ between rest-state (black) and stress-state (red) (two-way ANOVA: behaviour-state, F(1, 6195) = 60.67, p = 7.87×10^−15^; day, F(1, 6195) = 15.39, p = 8.84×10^−5^; behaviour-state x day, F(1, 6195) = 0.26, p = 0.61; Tukey’s HSD: acute-rest, n = 1625 cell pairs vs acute-stress, n = 1610 cell pairs, p = 1.2×10^−8^; chronic-rest, 1701 cell pairs vs chronic-stress, n = 1263 cell pairs, p = 3.8×10^−6^; acute-rest vs chronic-rest, p = 0.0016). **b** Relationship between cell pairs of positive co-activity Z-scores during both rest-state and stress-state significantly differs between days (acute: slope = 0.99, R^2^ = 0.022, p = 0.033 n = 208 cell pairs; chronic: slope = 1.74, R^2^ = 0.019, p = 0.076, n = 165 cell pairs; likelihood ratio test, p = 1.28×10^−7^) of CIS. Boxplots represent interquartile range (IQR, 25^th^-75^th^ percentiles), median is the thick line in the box and whiskers extend to 1.5 times the IQR. * p < 0.01, ** p < 0.005, *** p < 0.001. Acute-rest: N = 17 mice; acute-stress, N = 16 mice; chronic-rest, N = 16 mice; chronic-stress, N = 16 mice) .

## Discussion

Accumulating evidence has identified stress-induced changes in the hippocampus across biological scales – from behavioral deficits to their neuronal, synaptic and molecular correlates. However, how the intact hippocampal circuitry in an awake behaving animal responds to, and encodes information during, a stressful situation is poorly understood (McEwen, 2007). Here, we report a reduction in spiking in CA1 pyramidal neurons during the first (acute) exposure to 2-hour immobilization stress (Fig. 1d and e). However, multiple exposures to stress led to suppressed spiking even during rest periods outside of the stress itself. During both acute and chronic stress, theta/delta ratio remained low (Fig. 1g) and SPW-R events were of longer duration compared to rest (Fig. 2b and c). Further, stress altered the firing of CA1 pyramidal cells during SPW-Rs, leading to a greater fraction of spikes occurring inside SPW-Rs during both acute and chronic stress (Fig. 3c). Finally, pyramidal cells displayed enhanced co-activity which was also carried over to the rest-state on the final day of the 10-day chronic stress paradigm (Fig. 4a and b).

Considering that stress leads to elevated levels of glutamate and calcium, a long-held view in research on stress-induced plasticity is that stress leads to enhanced excitability in hippocampal circuits (Gunn and Baram, 2017; Sapolsky et al., 1986). However, our data from *in vivo* recordings revealed a robust decrease in firing rates of CA1 pyramidal cells during acute stress (Fig 1d). This result is consistent with a previous study that found suppression of pyramidal ‘place cell’ activity in an immobilized rat even when it was moved through its place field on a track (Foster et al., 1989) as well as other studies which reported suppressed place cell firing rates after the termination of stress (Park et al., 2015; Passecker et al., 2011). We also explored other measures of hippocampal activity by assessing pyramidal cell activity during SPW-Rs, which are high-frequency transient oscillations, referred to as “built in” sensors of hippocampal excitability (Buzsáki, 2015). Strikingly, this analysis revealed evidence in support of stress-induced enhanced excitability as pyramidal cells discharged a greater fraction of spikes inside SPW-Rs during stress states (Fig 3c). Together these results offer a new framework for analyzing stress-induced hyperexcitability at the network level, manifested as SPW-R events during stress.

Our analyses also revealed stress-induced enhancement in CA1 neuronal synchrony as evidenced by increased pyramidal cell co-activation during SPW-Rs (Fig. 4). Such co-activity during repeated exposures to stress is likely to cause CA1 pyramidal neurons to fire together, thereby creating conditions at the synaptic level that are consistent with the synaptic saturation hypothesis of stress (Diamond et al., 2004). This gives rise to questions that will require further investigation, including if such synchronous activity would saturate synaptic mechanisms that occlude further plasticity necessary for subsequent encoding? If so, this may offer insights into earlier findings that chronic stress causes rigidity in hippocampal networks (Tomar et al., 2015), thus leading to context generalization (Krugers et al., 1997) and spatial learning deficits (Kim et al., 2007; Park et al., 2015). Interestingly, a recent rodent study demonstrated a causal relationship between long duration SPW-Rs and learning (Fernández-Ruiz et al., 2019). In light of this study, our findings of increased pyramidal cell synchrony, along with long-duration SPW-Rs suggests that during stress, the hippocampus may engage in a learning state involved in encoding aversive memories related to the stressful experience (Cadle and Zoladz, 2015; Diamond et al., 2004, 2007; Schwabe et al., 2012).

The observation that altered SPW-R properties were present from the onset of immobilization stress (Fig. 2d and supplementary Fig. 2c) also raises interesting questions about underlying mechanisms. For instance, this timeline suggest that both altered SPW-R characteristics and suppressed hippocampal spiking during the acute stress are mediated by sympatho-adrenal medullary (SAM) system-activated neurochemicals, (Cadle and Zoladz, 2015; Gunn and Baram, 2017). Further, reports that norepinephrine application in hippocampal slices alters SPW-R properties (Haq et al., 2012) and suppresses CA1 pyramidal spiking (Bergles et al., 1996; Pang and Rose, 1987), suggests norepinephrine release as a likely mechanism contributing to the stress phenotypes observed in this study. Finally, our findings on suppressed CA1 spiking during the chronic-rest state is in agreement with the role of glucocorticoid receptor (GR) mediated suppression of hippocampal neuronal activity during stress (Joëls, 2018).

Following repeated stress, the enhanced pyramidal cell co-activation during SPW-Rs was also accompanied by participation of CA1 pyramidal cells in fewer SPW-Rs than during non-stress states (Fig. 3e and f). Whether suppressed participation of pyramidal cells is an epiphenomenon or is to counter enhanced network synchrony as stress becomes chronic, is not clear. However, these results show that repeated stress alters hippocampal ripple-spike interactions, a phenomenon previously linked to altered cognition in rodent models of disease (Middleton et al., 2018; Raveau et al., 2018; Suh et al., 2013; Witton et al., 2016). Two major afferents to area CA1 that influence information processing and SPW-R properties are the Schaffer collaterals from area CA3 (Davoudi and Foster, 2019; Nakashiba et al., 2009) and temporoammonic inputs from the entorhinal cortex (Oliva et al., 2018; Yamamoto and Tonegawa, 2017). We found no change in either SPW-R-associated low-gamma power (Fig. 2f) or ripple-burst patterns (Fig. 2g), suggesting that the stress phenotypes we observed are likely caused by changes occurring within areas CA1 and CA3 and are minimally influenced by entorhinal inputs. In addition, extrahippocampal regions, including the amygdala, are known to influence CA1 neural dynamics (Ghosh et al., 2013; Kim et al., 2012, 2015), LTP (Kim et al., 2005; Vouimba and Richter-Levin, 2005) and the release of stress hormones, including catecholamines (Roozendaal et al., 2009; Schwabe et al., 2012). Thus, future studies will be needed to characterize the relative contributions of CA3 and amygdalar inputs to SPW-R duration, pyramidal cell synchrony, and altered participation of pyramidal cells in SPW-Rs, reported in the current study.

In conclusion, a large body of earlier work has characterized the morphological, molecular, physiological and behavioral changes in the hippocampus following either acute or repeated stress. Our study adds a new layer of understanding by providing a window into the dynamics of hippocampal network activity during episodes of stress and identifies altered ripple-spike interactions as a potential biomarker of stress.

## Acknowledgements

We thank all members of the CBP laboratory for their support, the Advanced Manufacturing Support Team, RIKEN Center for Advanced Photonics for their assistance in microdrive production and Lalitha Krishnan for assistance with figure generation. We also thank Dr. Kazumasa Z. Tanaka for inputs and support. This works was supported by Grant-in-Aid for Scientific Research from MEXT (19H05646; T.J.M), Grant-in-Aid for Scientific Research on Innovative Areas from MEXT (19H05233; T.J.M) and RIKEN BSI and CBS (T.J.M).

## Author contributions

A.T. and T.J.M conceived the study. A.T. performed all experiments. A.T. D.P. and T.J.M. analysed the data. A.T., T.J.M. and S.C. wrote the manuscript with input from D.P. Funding provided by T.J.M.

## Correspondence and request for material

Any request for data or code should be addressed to A.T. or T.J.M.

## Methods

### Animals

All experiments were performed using male mice. In total 21 mice were used; of these, 4 were used for measuring the stress effects on bodyweight while the remaining 17 were used for in vivo electrophysiology. All mice were aged between 3 and 6 months at the start of experiments and were maintained on a 12-h light-dark cycle with *ad libitum* access to food and water. All procedures were approved by the RIKEN Institutional Animal Care and Use Committee and complied with all relevant ethical regulations.

### Experimental design and stress protocol

Mice underwent the same chronic immobilisation stress (CIS) protocol as described previously (Tomar et al., 2015). Briefly, mice experienced complete immobilization (2h/d, 10 consecutive days: Fig. 1a) in rodent immobilization bags, without access to food or water. All mice underwent the same experimental protocol as described previously, except 5 mice that experienced a familiar track on the first and 10th day of experiment followed by stress exposure. Here we examined the first 30 minutes of data recorded during CIS (stress-state) and the preceding quiescence period (rest-state). Recordings on the first day of CIS were termed ‘acute’ while those on the last day of experiment were termed ‘chronic’, providing four time points: i) acute-rest, ii) acute-stress, iii) chronic-rest and iv) chronic-stress. Rest-state data was recorded for ∼ 15-30 min and hence for temporal distributions, theta/delta ratio, correlation between theta/delta and SPW-R occurrence, the first 15 minutes of data was used.

### Surgery, recording and histology

Mice were anaesthetized using Avertin (2, 2, 2-tribromoethanol; Sigma-Aldrich, 476 mg/kg, i.p.) and were surgically implanted with a microdrive (manufactured with the assistance of the Advanced Manufacturing Support Team, RIKEN Center for Advanced Photonics, Japan). The microdrive housed eight independently movable tetrodes (14 µm diameter, nichrome) and was placed above right dorsal hippocampus (coordinates from bregma: AP -1.8mm; ML + (1.2mm). Prior to surgery, tetrodes were gold plated to lower impedance down to a range of 100–250 kΩ. Tetrodes were gradually lowered over the course of several days, such that by the start of the experiment they reached the CA1 stratum pyramidale. Data were acquired using a 32-channel Digital Lynx 4SX acquisition system (Neuralynx, Bozeman, MT). Signals were sampled at 32,556 Hz and spike waveforms were filtered between 600 Hz and 6 kHz. Skull screws located above the cerebellum served as a ground, and a tetrode that was seated in the superficial layers of the neocortex, and devoid of spiking activity, was used for referencing. 3-4 weeks after surgery, when all tetrodes reached the CA1 stratum pyramidale, evident by multiple large amplitude spikes and SPW-Rs, the experiment was initiated. During both rest-state and stress-state recordings the mice were located in a small circular sleep box (15 cm diameter). At the conclusion of the experiment mice underwent terminal anaesthesia (Avertin), and electric current (30 µA, for 8 s) was administered through each electrode to mark their locations. Transcardial perfusion was carried by using saline followed by 4% paraformaldehyde (PFA) followed by a further 24 h fixation in 4% PFA. Brains were sliced using a vibratome (Leica) to prepare coronal slices (50 µm thick), which were subsequently stained with Eosin Y, and inspected by standard light microscopy to confirm electrode placement.

### Unit isolation and spike analysis

Spike sorting was performed by automatic spike sorting program (KlustaKwik; (Harris et al., 2000)), followed by manual adjustments of the cluster boundaries using SpikeSort3D software (Neuralynx). Candidate clusters with < 0.5% of spikes displaying an inter-spike-interval shorter than 2 ms, a total number of spikes exceeding 50, having a cluster isolation distance value (Schmitzer-Torbert et al., 2005) >15, spike width (peak-to-trough) >170 µs and complex spike index (CSI) (McHugh et al., 1996) >5 were considered as pyramidal cells and were used for further analysis.

### Local field potential analysis

The raw LFP data were first downsampled using a custom software written in C to 1627.8 Hz (a factor of 20), followed by a quality control measure and channel selection via visual inspection and biggest power in SPW-R frequency band (80-250 Hz). A low-pass filter with a cut-off frequency equal to half the target sampling frequency was applied to the LFP prior to downsampling to prevent any signal distortion. Power spectral density (PSD) was calculated by using Welch’s averaged modified periodogram method (pwelch function in Matlab) with a 2048 sample window size (1.26 s), 50% overlap and 4096 FFT points (2.52 s) resulting in a time-varying spectrogram. To account for power fluctuations caused by difference in position/impedance of the electrodes and to make PSD values comparable across mice, each PSD curve was normalized by its own mean power within the 0 – 3 Hz frequency band. Temporal dynamics between behaviour states were assessed by binning LFP data in 1 min time-bins and by comparing this data between stress-states and rest-states.

### SPW-R detection

SPW-R events were detected using modifications to the method described in (Csicsvari et al., 1999b). Previously selected LFP channels were first band-pass filtered (80-250 Hz) using a 69 order Kaiser-window FIR zero-phase shift filter. Subsequently, the absolute value of Hilbert transform (instantaneous ripple power) was smoothed with 50 ms Gaussian window and candidate SWR-events were detected as periods where magnitude exceeded 3 standard deviation (SD) above its mean for longer than 30 ms. Further, the initiation and termination points of candidate SWR events were defined as points when the magnitude returned to the mean. Summed multi-unit activity (MUA) across all pyramidal cells fired during any given recorded session was converted to instantaneous firing rate using time bin size equal to the underlying LFP trace, sampling rate and smoothed, to allow detection of firing bursts using the same thresholds as described for the candidate SPW-R events detection. Candidate SPW-R events not coincident with MUA bursts were excluded from subsequent analysis. For precise peak PSD frequency detection of the SPW-R events, a multitaper method was employed on the product of each filtered SPW-R waveform and Hanning window of the same length.

### Single Unit properties

Mean firing rate of individual single units was calculated as total number of spikes emitted by the unit during sleep/stress trial divided by the trial’s duration. Peak firing rate of each single unit was calculated by smoothing ISIs of the unit with a 5SD Gaussian kernel and taking maximum value of the resulting firing rate curve (firing rate over time). Spike bursting analysis was performed as described previously (Bakkum et al., 2014) by defining two or more spikes occurring within 10 ms time bin as a burst.

### SPW-R triggered spectrograms

The spectrograms were calculated by using complex wavelet transform (CWT) (Morlet type wavelet, parameter = 7) method applied to a segment of the wide band LFP in 400 ms window centred on each SPW-R. Resulting spectrograms were averaged across individual SPW-R events in each recording session/trial. To compensate for 1/f power loss each power value within each spectrogram was multiplied with the frequency correspondent to that power value. To make power values comparable across subjects, each spectrogram was normalized by its own mean power within (0 – 5 Hz) frequency band.

### SPW-R bursting analysis

The concentration of SPW-R events in time (bursting) was estimated by first taking middle time stamp of each SPW-R event and calculating the difference between time stamps of the previous and next event. If adjacent events occurred further than 200ms apart the current event marked as ‘singlet’. Number of timestamps in remaining events was counted, and each event marked as member of ‘doublet’, ‘triplet’ or ‘other’ (for more than 3 events in a SPW-R burst).

### Participation of single units in SPW-R events

Each ripple event’s start and end timestamps were used to quantify single unit activity corresponding to the co-occurred SPW-R events. Within SPW-R firing rate of any given single unit within each trial was calculated as total number of spikes generated by the single unit divided by combined duration of all SPW-R events of the trial. Between SPW-R firing rate for any given trial was calculated by first, removing all spikes fired within all SPW-R events occurred during the trial and then calculating mean firing rate of resulting spike train. Participation of a single unit in SPW-R events was calculated as % of SPW-R events the unit fired at least single spike in any given trial.

### Coactivity Z-score

A likelihood of any given pair of single units (limited to pyramidal cells) firing together during SPW-R events or “coactivity Z-score” was calculated as described previously (Singer and Frank, 2009). Briefly, for any given trial a set of start and stop timestamps of SPW-R events and a set of spike train timestamps, fired by pyramidal cells were prepared. Then a ‘coactivity matrix’ of size (Ncells x Nspw-r events) was calculated. Each element of the coactivity matrix is set to 1 if a given cell was active (e.g. fired at least one spike) during given SPW-R event or 0 otherwise. Next, for every possible pair of cells a raw coactivity score was calculated as number of SPW-R events during which both cells of the pair were active. Finally, a z-cored coactivity value was calculated by normalisation of difference between raw coactivity score and variance by standard deviation as described in (Singer and Frank, 2009).

### Statistical analysis

All statistical analyses were performed in R software (3.3.2). All boxplots were analysed using two-way ANOVAs (*aov* function, stats package) followed by Tukey’s honestly significant difference (HSD) test. (*TukeyHSD* function, stats package). Distributions of various SPW-R properties as a function of SPW-R durations/amplitude were assessed using linear mixed effects models (LMMs) where mouse identity was specified as a random factor and behaviour states and categorical variables were specified as fixed factors (*lmer* function, lme4 package). The output of the *lmer* function was summarized as an ANOVA table (*anova* function, stats package). Post hoc pair-wise comparisons were made using the least-squares-means (LSM) approach (*lsmeans* function, lsmeans package). Correlation between parameters was calculated using Pearson’s correlation coefficient analysis (base package). Dependence of a parameter on another was calculated by employing standardized major axis (SMA) regression (*sma* function, smatr package). For the log-transformed scatter plots, the slope and the R^2^ values were calculated using log-transformed data and accordingly reported. Comparisons between regression lines were made by likelihood ratio tests (*sma* function, smatr package). For cumulative distribution analysis, the Kolmogorov-Smirnov test was employed (*ks*.*test*, stats package). Boxplots represent Interquartile Range (IQR, 25^th^-75^th^ percentiles), median is the thick line housed in the box and whiskers extend to 1.5 times the IQR. No data point was removed as outlier either for making boxplots or for statistical analysis. Unless noted, the level of statistical significance was set to 0.05 and p values are shown as follows: * p < 0.05; ** p < 0.01; *** p < 0.001.

### Data and code availability

The data and MATLAB codes that support the findings of this study are available from corresponding authors upon request.

---

**Supplementary Fig. 1.**
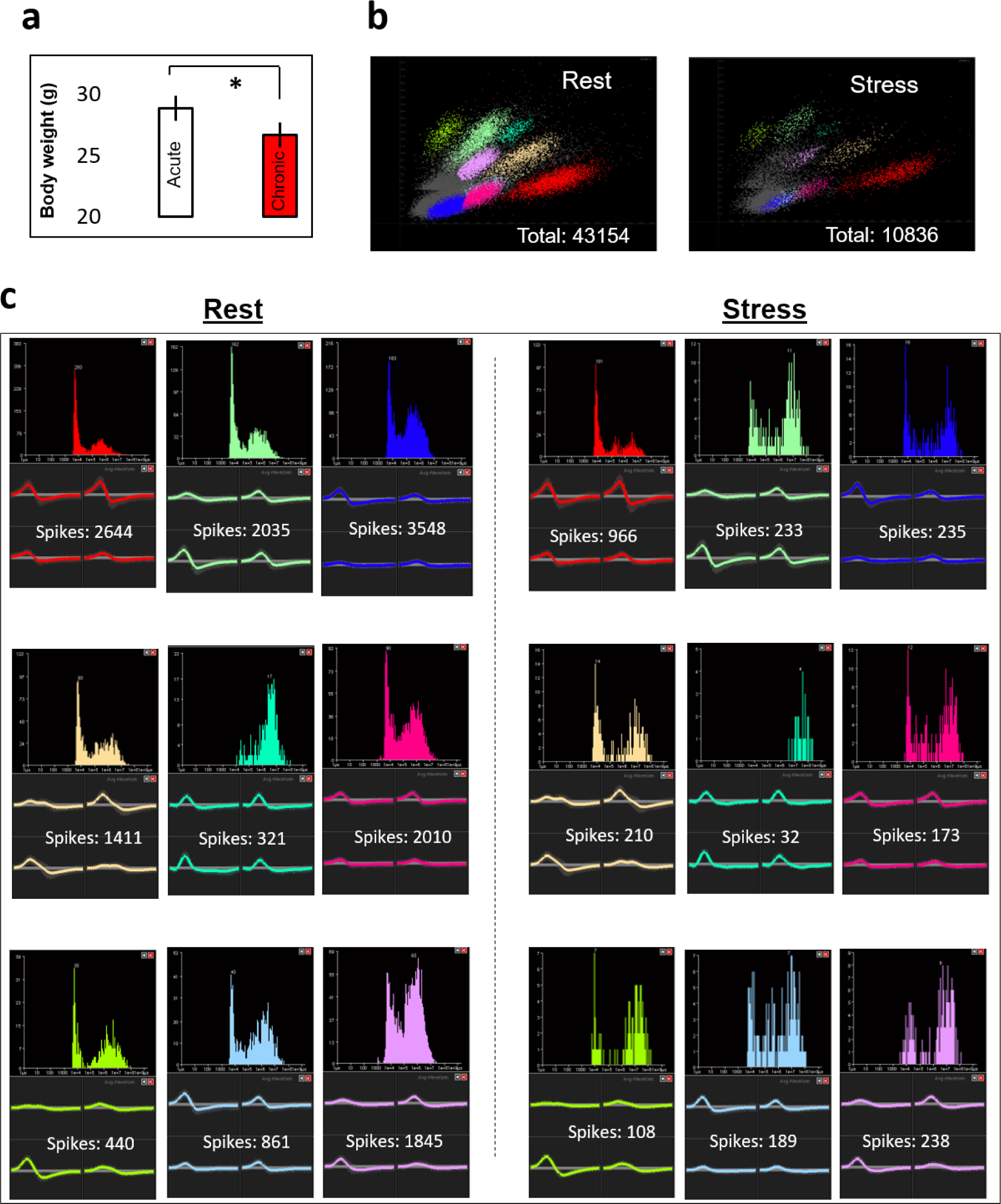
**a** Change in absolute body weight with the progression of CIS protocol (n = 4 mice). **b** Time matched (30 min) representative examples of unsorted spikes recorded during rest-state (left) and stress-state (right). Numbers next to each cluster depict total number of spikes recorded during that state. **c**. Each subpanel during rest-state (left) and stress-state (right) contains inter-spike-intervals (top) and waveforms (bottom) on each wire of a tetrode from all 9 cells sorted in b. Numbers on each subpanel represents total number of spikes discharged by that specific cell. * p < 0.05.

**Supplementary Fig. 2.**
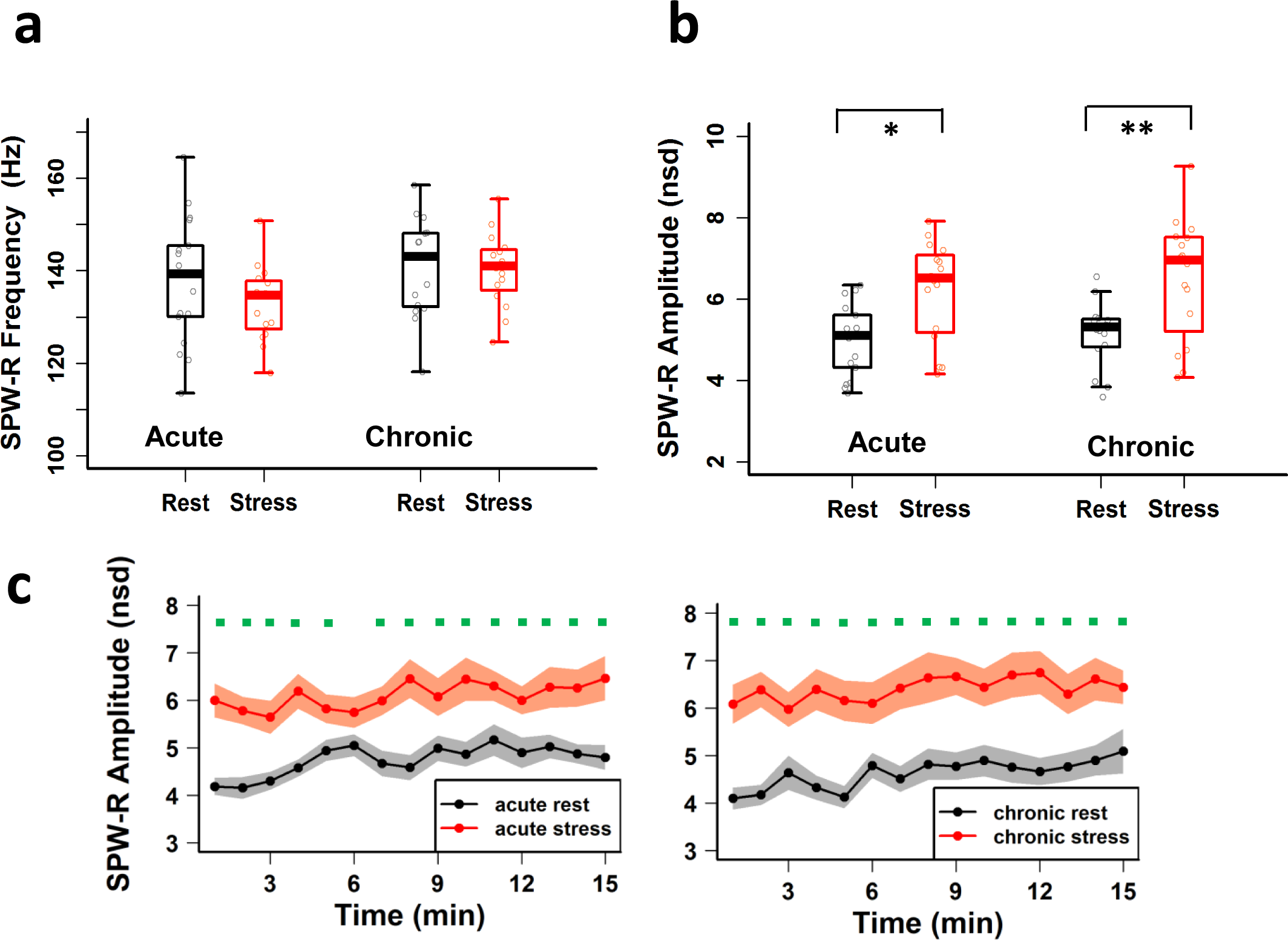
**a** Frequency of SPW-R oscillations did not differ between rest-state and stress-state (two-way ANOVA: behaviour-state, F(1, 61) = 1.106, p = 0.297; day, F(1, 61) = 3.838, p = 0.055; behaviour-state x day, F(1, 61) = 0.687, p = 0.41). **b** SPW-R amplitude differs between rest-state and stress-state (acute-rest vs acute-stress, p = 0.013; chronic-rest vs chronic-stress, p = 0.0066). **c** Temporal dynamics of averaged SPW-R amplitudes (1-min bins) differ between behaviour states on first day (left: LMMs: behaviour-state, F(1, 447) = 237.47 p < 2.22×10^−16^; minutes, F(14, 447) = 1.91, p = 0.023; behaviour-state x minutes, F(14, 447) = 1.01, p = 0.44) and the last day (right: LMMs: behaviour-state, F(1, 412) = 244.57, p < 2.22×10^−16^; minutes, F(14, 412) = 1.904, p = 0.024; behaviour-state x minutes, F(14, 412) = 0.696, p = 0.78) of CIS. Green dots on the top of graph indicate significant differences between behaviour-states. The boxes in the box plots, represent interquartile range (IQR, 25^th^-75^th^ percentiles), median is the thick line in the box and whiskers extend to 1.5 times the IQR. * p < 0.05, ** p < 0.01, *** p < 0.001. Acute-rest, N = 17 mice; acute-stress, N = 16 mice; chronic stress, N = 16 mice; chronic stress, N = 16 mice).

**Supplementary Fig. 3.**
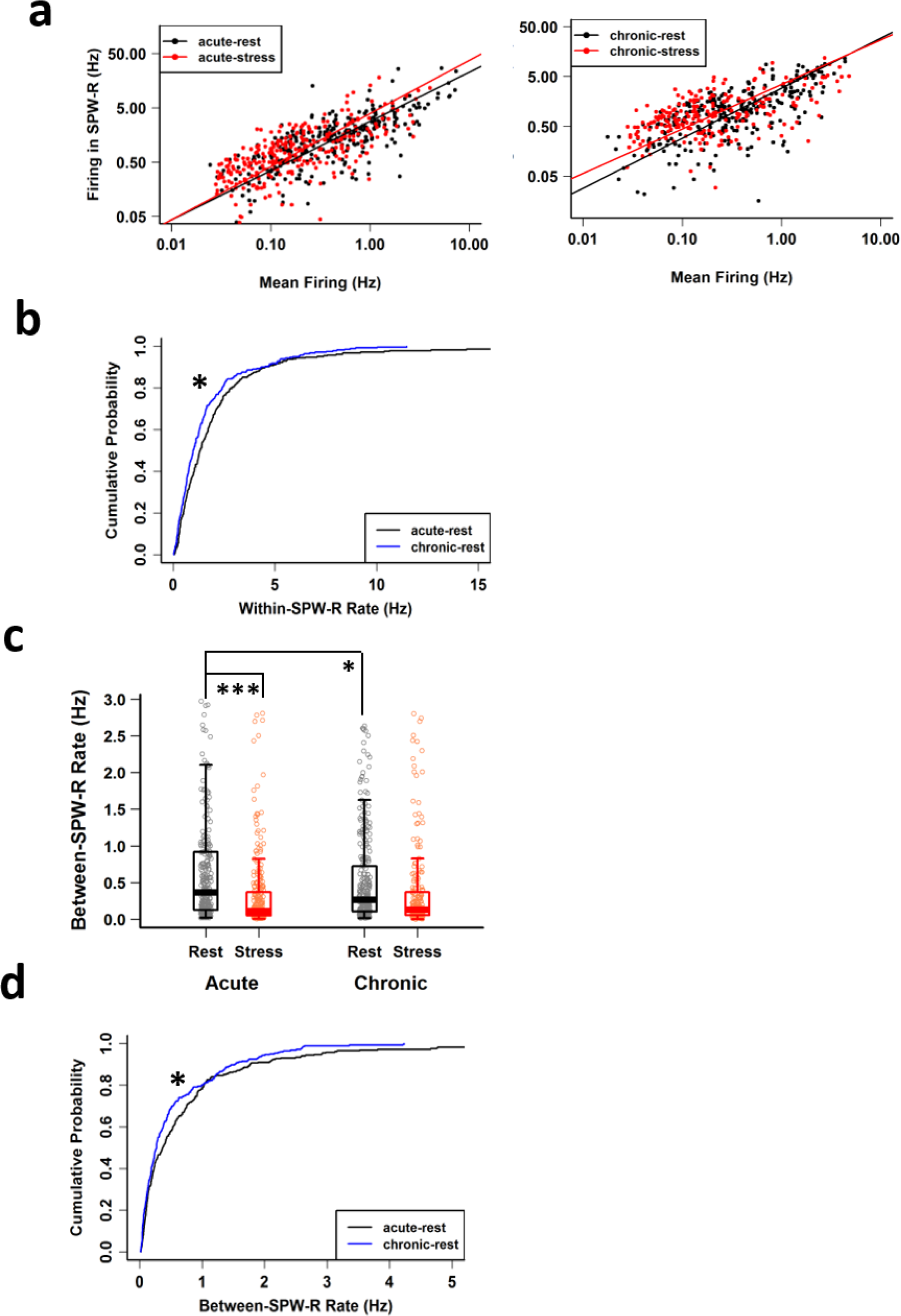
**a** Dependency of CA1 pyramidal cell firing rate in SPW-Rs on their mean firing rate was not affected by either acute (acute-rest: slope = 0.907, R^2^ = 0.56, p < 2.22×10^−16^; acute-stress: slope = 0.92, R^2^ = 0.47, p < 2.22x 10^−16^; likelihood ratio test (df = 1) = 0.063, p = 0.80) or chronic stress (chronic-rest: slope = 1.003, R^2^ = 0.44, p < 2.22×10^−16^; chronic-stress: slope = 0.89, R^2^ = 0.27, p < 2.22×10^−16^; likelihood ratio test (df = 1) = 2.62, p = 0.105). **b** Cumulative distribution plots for within SPW-R firing rate differ between rest-states on the first (black line) and the last (blue line) day of CIS (acute-rest vs chronic-rest: Kolmogorov-Smirnov test, D = 0.133, p = 0.013). **c** Between SPW-R average firing rate differs between behaviour states (two-way ANOVA: behaviour-state, F(1,1074) = 31.042, p = 3.191 ×10^−8^; day, F(1,1074) = 0.644, p = 0.422; behaviour-state x day, F(1,1074) = 9.428, p = 0.0022; Tukey’s HSD: acute-rest vs acute-stress, p = 6.01 ×10^−9^; acute-rest, vs chronic-rest, p = 0.021; chronic-rest vs chronic-stress, p = 0.394). **d** Cumulative distribution plots of between SPW-R firing rate differ between rest-states on the first (black line) and the last (blue line) day of CIS (acute-rest vs chronic-rest: Kolmogorov-Smirnov test, D = 0.119, p = 0.036). The boxes in the box plots, represent interquartile range (IQR, 25^th^-75^th^ percentiles), median is the thick line in the box and whiskers extend to 1.5 times the IQR. * p < 0.05, ** p < 0.01. Acute-rest: n = 288 cells, N = 17 mice; acute-stress: n = 282 cells, N = 16 mice; chronic-stress: n = 282 cells, N = 16 mice; chronic-stress: n = 226 cells, N = 16 mice.

